# Immunodominant and neutralizing linear B cell epitopes spanning the spike and membrane proteins of Porcine Epidemic Diarrhea Virus

**DOI:** 10.1101/2021.10.05.463270

**Authors:** Kanokporn Polyiam, Marasri Ruengjitchatchawalya, Phenjun Mekvichitsaeng, Kampon Kaeoket, Tawatchai Hoonsuwan, Pichai Joiphaeng, Yaowaluck Maprang Roshorm

## Abstract

Porcine Epidemic Diarrhea Virus (PEDV) is the causative agent of PED, an enteric disease that causes high mortality rates in piglets. PEDV is an alphacoronavirus that has high genetic diversity. Insights into neutralizing B cell epitopes of all genetically diverse PEDV strains are of importance, particularly for designing a vaccine that can provide broad protection against PEDV. In this work, we aimed to explore the landscape of linear B cell epitopes on the spike (S) and membrane (M) proteins of global PEDV strains. All amino acid sequences of the PEDV S and M proteins were retrieved from the NCBI database and grouped. Immunoinformatics-based methods were next developed and used to identify putative linear B cell epitopes from 14 and 5 consensus sequences generated from distinct groups of the S and M proteins, respectively. ELISA testing predicted peptides with PEDV-positive sera revealed 9 novel immunodominant epitopes on the S protein. Importantly, 7 of these novel immunodominant epitopes and other subdominant epitopes were demonstrated to be neutralizing epitopes by neutralization-inhibition assay. Additionally, our study shows the first time that M protein is also the target of neutralizing antibodies as 7 neutralizing epitopes in the M protein were identified. Conservancy analysis revealed that epitopes in the S1 subunit are more variable than those in the S2 subunit and M protein. In this study, we offer the immunoinformatics approach for linear B cell epitope identification and a more complete profile of linear B cell epitopes across the PEDV S and M proteins, which may contribute to the development of a greater PEDV vaccine as well as peptide-based immunoassays.

## INTRODUCTION

A swine disease named Porcine Epidemic Diarrhea (PED) is one of the main diseases that causes huge economic losses in swine industry worldwide, including Asia, America and Europe (1, 2). Porcine Epidemic Diarrhea Virus (PEDV) is the causative agent of PED (3). PEDV can infect and cause symptoms in all ages of pigs; however, it has more effect on suckling piglets with 80-100% death rate (4). PEDV is a member of the *Alphacoronavirus* genus and it is an enveloped, positive-sense, single-stranded RNA virus (2). PEDV genome is approximately 28 kb long that encodes three nonstructural proteins and four structural proteins, which are spike protein (S), membrane protein (M), envelope protein (E), and Nucleocapsid protein (N) (2).

Studies on the genetic profile of PEDV have demonstrated that the PEDV genome is highly diverse (5). Outbreaks and re-emergences of PEDV with high genetic diversity have been reported in many countries (6). Even though vaccination is an effective method to prevent and control infection, the high genetic variation of PEDV remains one of the challenges in designing an effective PEDV vaccine. Although both B cells and T cells can be elicited by vaccines, it is generally though that most vaccines confer protection through the induction of B cells to produce neutralizing antibodies (7). Hence, understandings of antibody responses following PEDV infection in regard to the landscape of immunodominant and neutralizing B cell epitopes as well as conserved and unique epitopes in distinct strains will facilitate designing a more powerful universal vaccine that can cope with all diverse strains of PEDV.

While M protein is the most abundant in the PEDV particle and plays an important role in the viral assembly process (8, 9), the PEDV S protein interacts with the host receptor and is composed of immunogenic regions capable of inducing neutralizing antibodies (10, 11). The S protein is thus considered the main target for vaccination. Similar to other coronaviruses, the PEDV S protein is a large glycoprotein composed of 1383 amino acids (based on the classical strain PEDV CV777) and it can be divided into 2 functional subunits: (i) the N-terminal S1 subunit, responsible for receptor binding, and the C-terminal S2 subunit, responsible for membrane fusion (12, 13). The S1 subunit is comprised of N-terminal domain (NTD) and the CO-26K equivalent (COE) domain (residues 499-638), which is responsible for receptor binding (12, 14). The S2 subunit consists of 3 domains: a large ectodomain, a transmembrane domain and a cytoplasmic tail or endodomain (12, 15). A large ectodomain is composed of protease cleavage site, fusion peptide (FP), 2 heptad repeat (HR1 and HR2) regions, which play important roles in viral and host cell membrane fusion (12, 15, 16).

The COE domain has been identified as an immunogenic domain containing B cell epitopes recognized by neutralizing antibodies (11, 14, 17); therefore, it is considered an alternative vaccine target and has been extensively used in the development of recombinant PEDV vaccines (18–20). In the M protein, an epitope named M-14 has been identified from the PEDV CH/SHH/06 strain (8). In the S protein, 4 linear B cell epitopes including (i) S1D5 with SS2 as a core epitope (21), (ii) S1D6 with SS6 as a core epitope (21), (iii) peptide M (22), and (iv) 2C10, a neutralizing B cell epitope located at the C-terminus of the S protein (23, 24), were first identified. More recently, two conformational neutralizing epitopes located in the S1 NTD and COE were identified from truncated S proteins of the PEDV PT strain (11). Additionally, neutralizing epitopes located at the same region with the S1D5 and S1D6 epitopes were reported (10, 21). All these epitopes were identified based on experimental methods such as ELISA with truncated proteins, pepscan, and phage display, which are laborious, costly, and time-consuming. Additionally, by using these techniques, B cell epitope identification can focus only on some regions of the proteins, while a complete profile of B cell epitope across the entire proteins are of importance for vaccine design.

Recently, immunoinformatics has been demonstrated to be a powerful tool for identification of B and T cell epitopes (25, 26) as well as for vaccine design and *in silico* evaluation (27, 28). Importantly, immunoinformatics approach can facilitate big data analysis with less cost and time compared to experimental methods. Currently, there are many tools available for *in silico* prediction of B cell epitope. Among the epitope prediction resources, IEDB database provides multiple tools for B cell epitope prediction with high accuracy (29). Generally, the peptides predicted by immunoinformatics methods cannot be claimed as epitopes, although they are well characterized as potential B cell epitopes. Thus, experimental validation is crucial and necessary for confirming whether predicted epitopes are genuine epitopes. Based on this combined method, a complete set of B cell epitopes across the entire protein can be identified.

In the present work, we aimed to identify linear B cell epitopes of two PEDV structural proteins, S and M, from all diverse strains available in the database and to characterize their potential as neutralizing epitope. The amino acid sequences of the PEDV S and M available in the NCBI database were retrieved and phylogenetic tree were generated. Consensus sequences obtained from each group were then subject to linear B cell epitope prediction using multiple immunoinformatics tools. We generated our own methods for selecting potential B cell epitopes based on 4 known epitopes (SS2, SS6, peptide M, and 2C10). The selected peptides were then experimentally validated using ELISA by testing synthetic peptides with PEDV-positive sera. Their potential as a neutralizing epitope was then investigated using neutralization-inhibition assay. Based on this approach, the whole set of neutralizing linear B cell epitopes as well as immunodominant epitopes across the PEDV S and M protein were defined.

## RESULTS

### The S and M proteins from global PEDV strains are classified into 14 and 5 groups

Amino acid sequences of the PEDV S and M proteins were retrieved from all sequences in the NCBI database. After filtering with Sublime Text 3 to remove incomplete and repeated sequences, 560 and 924 sequences of the M and S proteins, respectively, were obtained. The sequences of each protein were next grouped and phylogenetic trees were generated. Sequences of the M protein were classified into 5 groups, while sequences of the S protein were classified into 14 groups (**Fig. 1**). The consensus sequence of each group of both proteins were next generated and aligned as shown in **Supplementary Fig. 1** and **2**. Percent similarity among different groups was in the range from 32 to 99.71% for the S protein and from 20.35 to 99.55% for the M protein.

**Figure 1.**
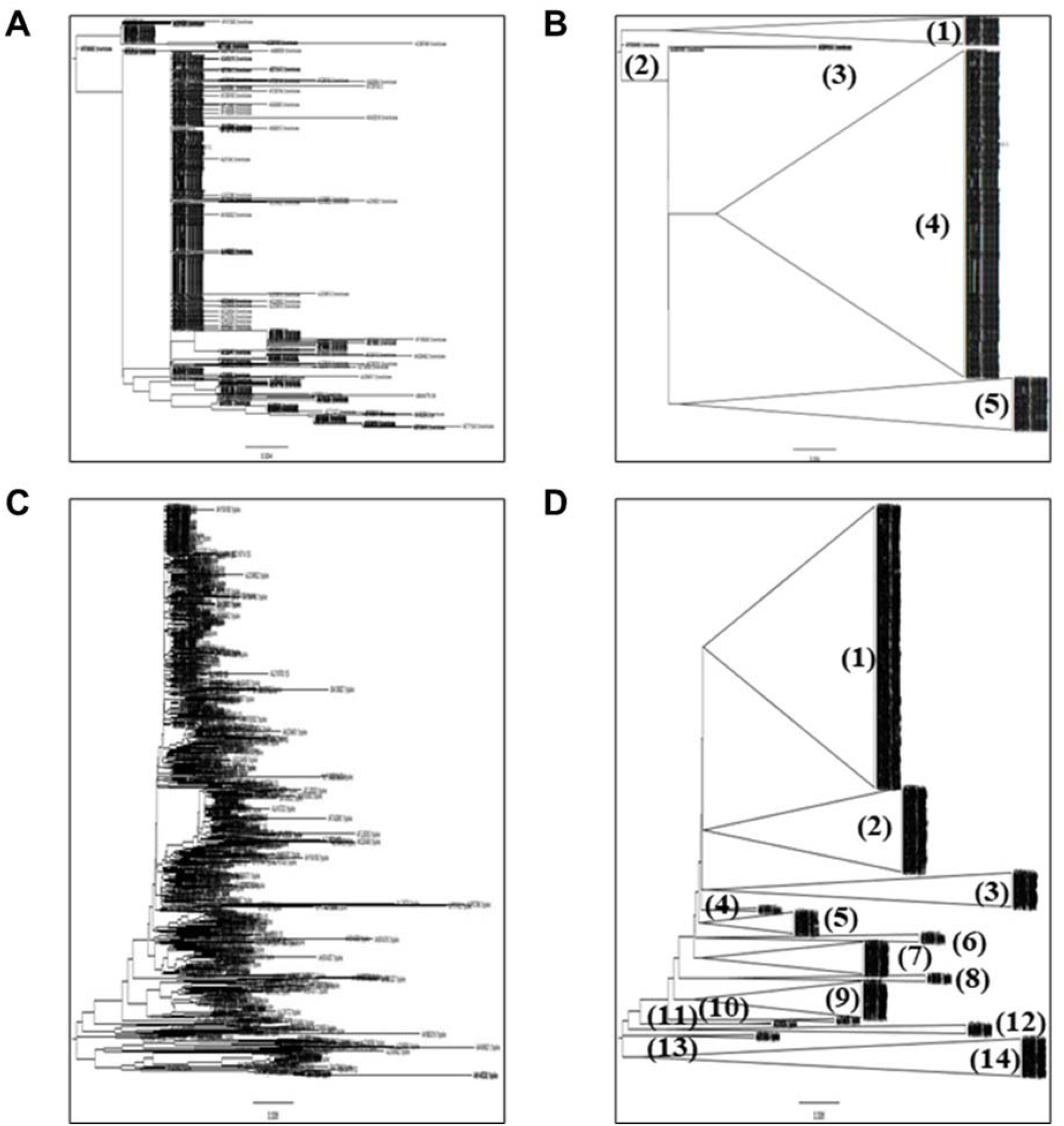
Phylogenetic tree of the PEDV M and S proteins. **(A** and **C)** Original phylogenetic tree of M and S proteins generated by SeaView program. (**B** and **D)** Groups of the M and S proteins generated by FigTree tool. Number represents group’s number.

### Three bioinformatics approaches are developed and used to predict and select linear B cell epitopes

We employed immunoinformatics to screen for B cell epitopes from diverse PEDV strains. Linear B cell epitopes have been suggested to be correlated with surface accessibility, coil probability, antigenicity, and hydrophilicity (30, 31); thus, we utilized multiple immunoinformatics tools to predict linear B cell epitopes and peptide characteristics. BepiPred-2.0 was used to predict linear B cell epitopes. In parallel, other features of the peptides were predicted using IUPred as well as the methods of Emini. Kolaskar & Tongaonkar, and Parker as shown in **Fig. 2B.**

**Figure 2.**
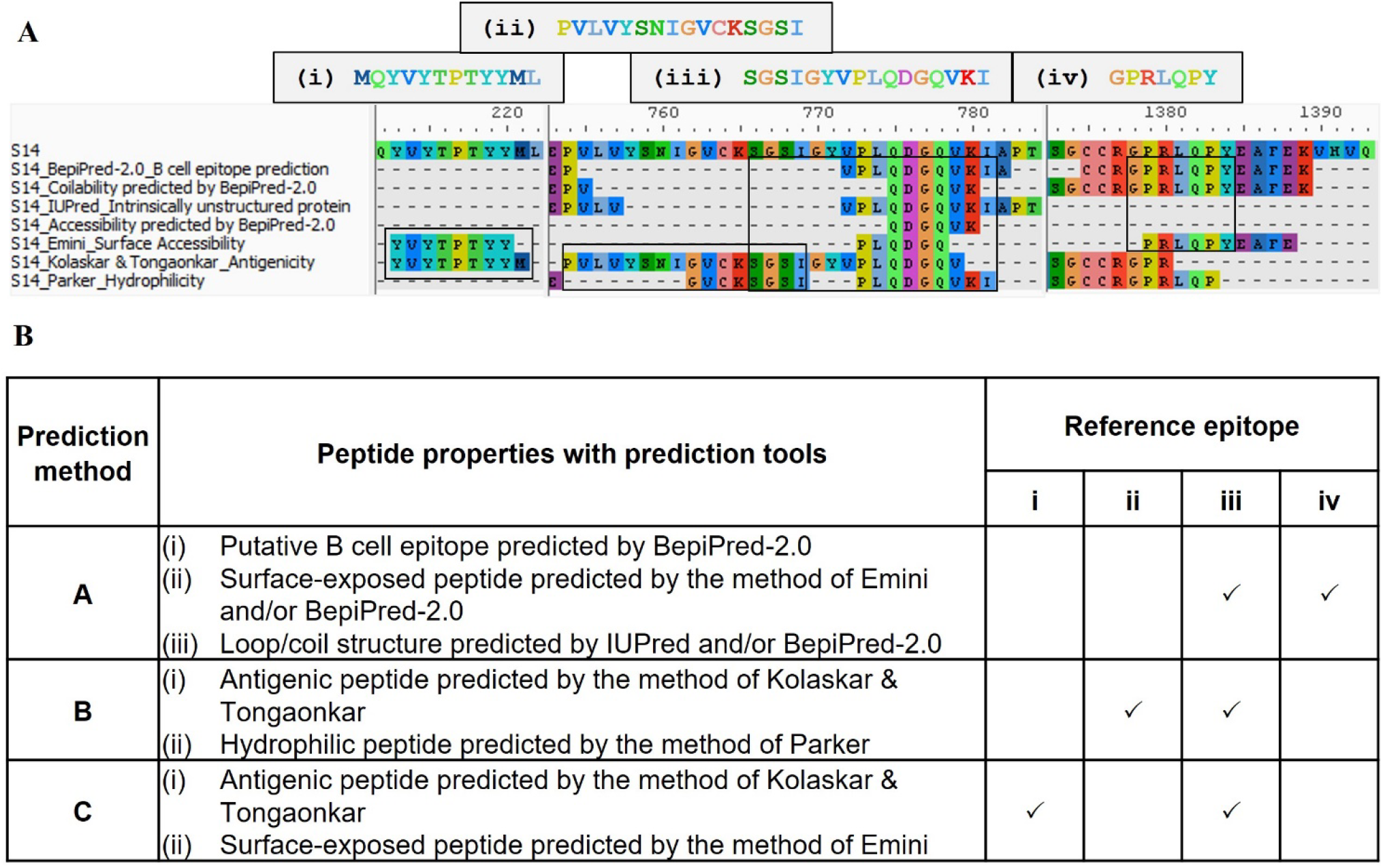
Immunoinformatics prediction and methods for B cell epitopes identification. **(A)** Immunoinformatics prediction and mapping of the known epitopes to the peptides predicted by different prediction tools. Predicted peptides obtained from each tool were mapped to 4 published epitopes. **(B)** Methods for identification of B cell epitopes. Three methods (A, B and C) were generated based on mapping results of the 4 published epitopes to the peptides predicted by immunoinformatics tools.

To assure that all important epitopes are picked up, we developed our own methods to identify and select putative linear B cell epitopes. Firstly, we aligned all predicted peptides obtained from each prediction tool. Four well known B cell epitopes of the PEDV S protein including (i) peptide M (MQYVYEPTYYML) (22), (ii) S1D5 (PVLVYSNIGVCKSGSI) (21), (iii) S1D6 (SGSIGYVPLQDGQVKI) (21) and (iv) 2C10 (GPRLQPY) (23, 24), were exploited as reference epitopes (**Fig. 2A**). Based on sequence alignment, we then created 3 different methods, designated methods A, B and C, for identifying and selecting linear B cell epitopes as described in **Fig. 2B.**

### Potential linear B cell epitopes are identified in 33 and 7 regions of the PEDV M and S proteins

The consensus sequences from all 14 groups of the S protein and 5 groups of the M protein were subject to linear B cell epitope prediction. Putative B cell epitopes were identified and selected based on methods A, B and C (**Supplementary Fig. 3** and **4)**. Note that the predicted peptides that overlap or are located in close proximity were combined into one lone peptide. Prediction of the M protein yielded 7 putative B cell epitopes, designated M-1 to M-7 (**Fig. 3** and **Table 1**). A total of 50 peptides in 33 regions of the S protein, designated S1B and S2B for the epitopes located on the S1 and S2 subunits, respectively, were predicted as B cell epitopes (**Fig. 4** and **Table 1**). In the COE region, 3 epitopes, namely S1B-15, S1B-16 and S1B-17, were identified. Most of the B cell epitopes predicted from different consensus sequences are located on the same regions. Impressively, our immunoinformatics methods could identify all published epitopes in the S protein as indicated in **Table 1** and **Supplementary Figure 5** and **6**.

**Table 1.**
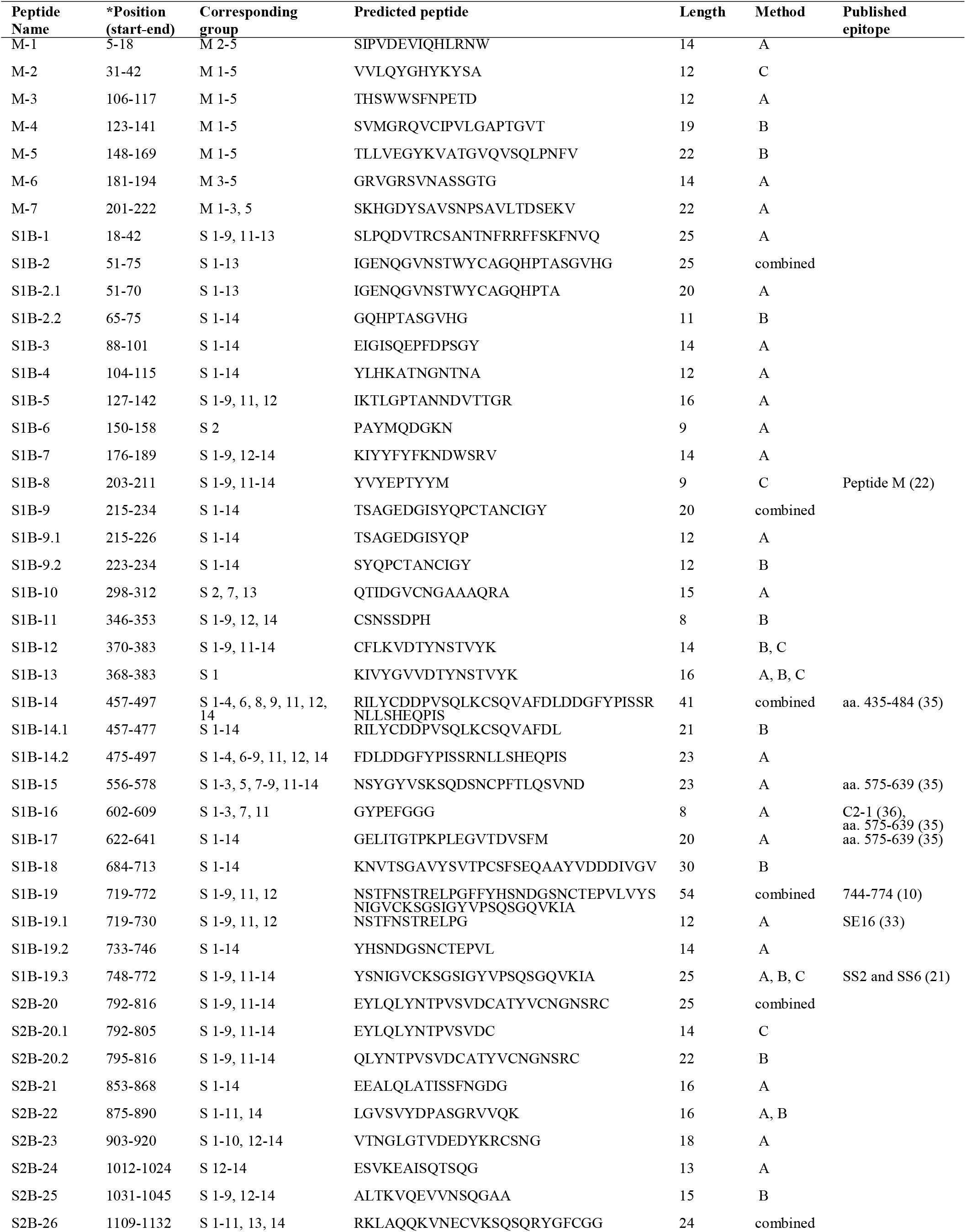
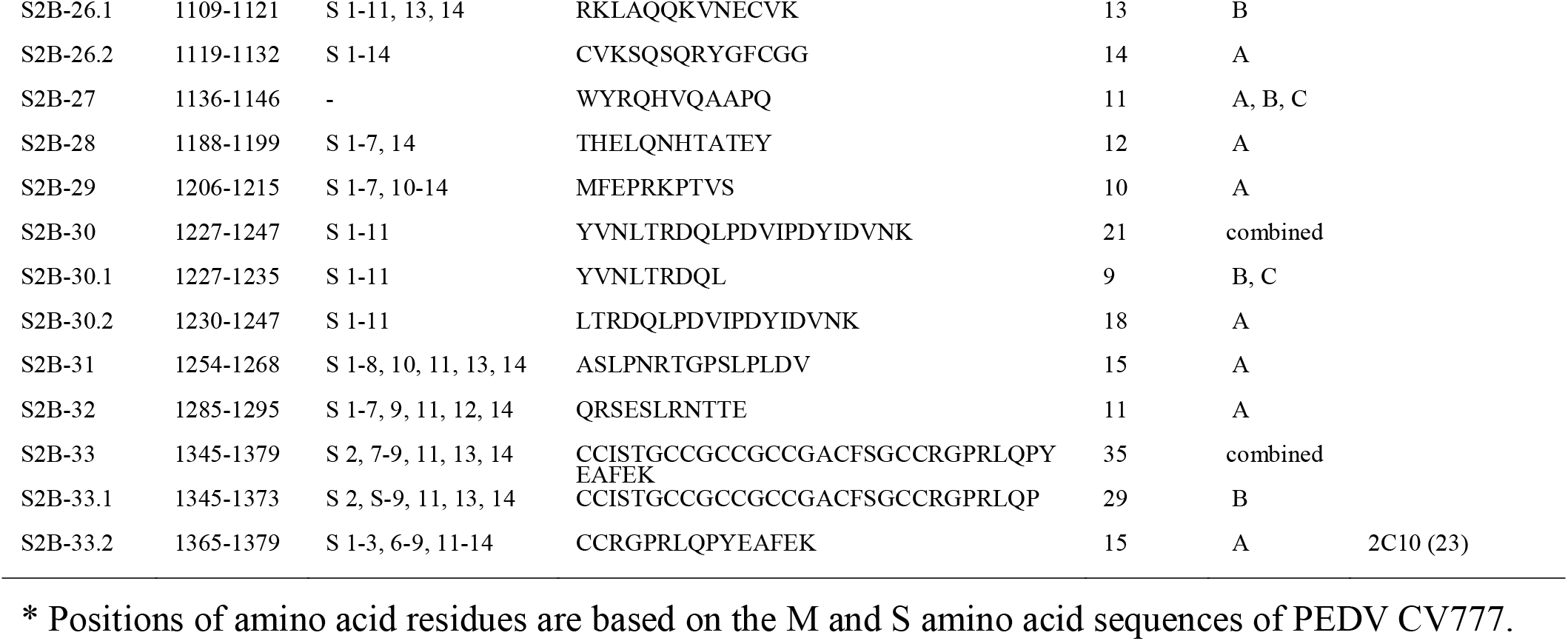
Predicted B cell epitopes of the PEDV M and S proteins that were selected for further experimental validation.

**Figure 3.**
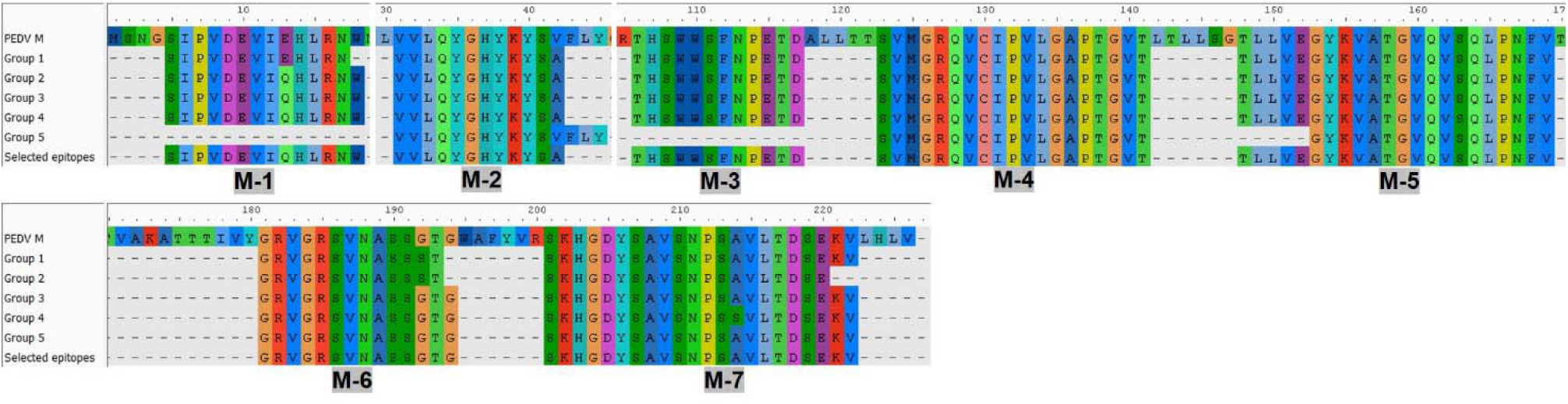
Predicted B cell epitopes of the M protein. B cell epitopes were predicted from consensus sequences of 5 groups of the M protein. Epitopes resulting from prediction of each group are shown and epitopes selected for further experimental evaluation are indicated in the “selected epitope” line with designated name.

**Figure 4.**
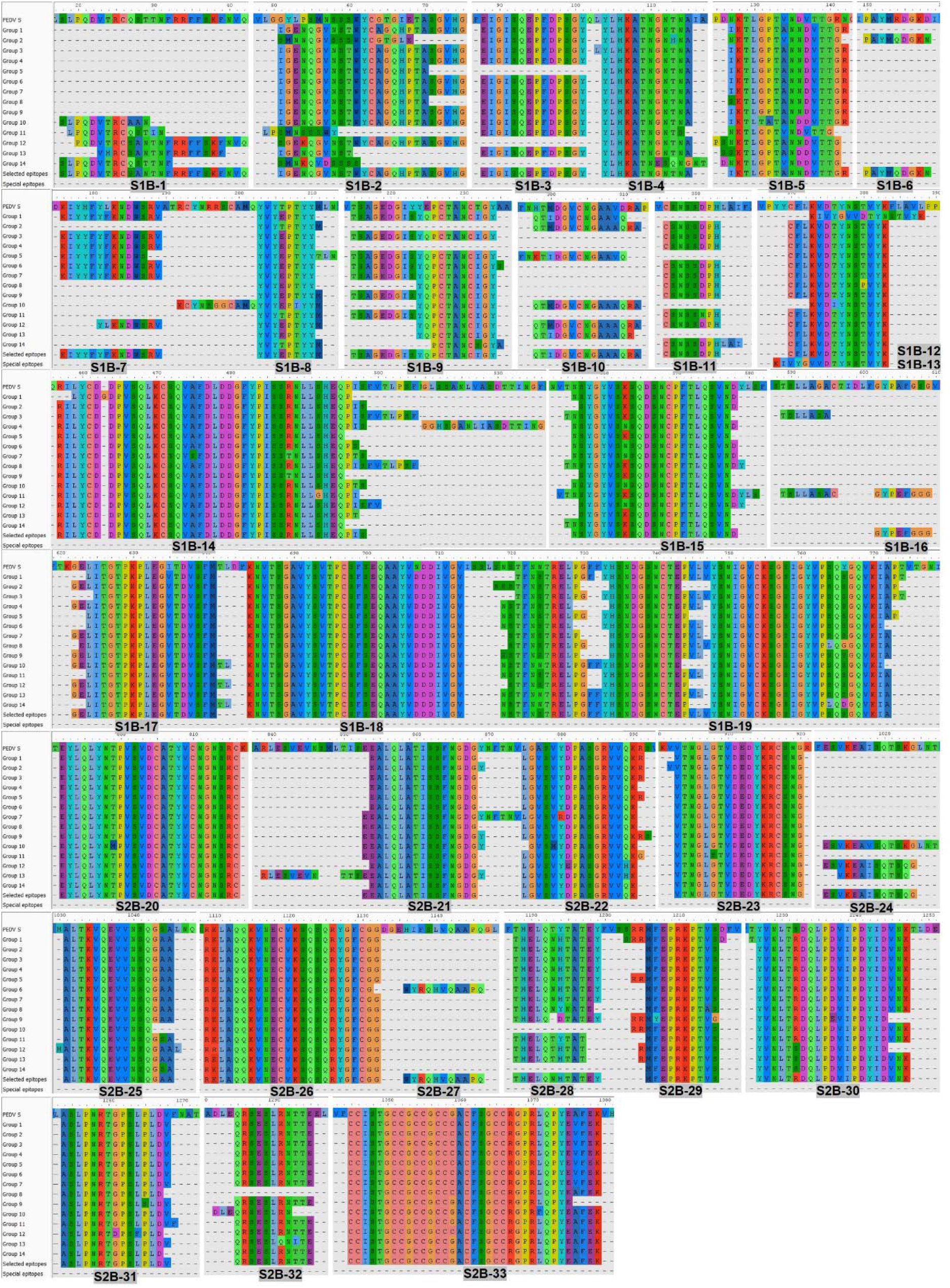
Predicted B cell epitopes of the PEDV S protein. B cell epitope prediction was carried out using consensus sequences derived from 14 groups of the S protein. Predicted epitopes from each group are shown. The epitopes selected for further experimental evaluation are indicated in the “selected epitope” line with designated name.

**Figure 5.**
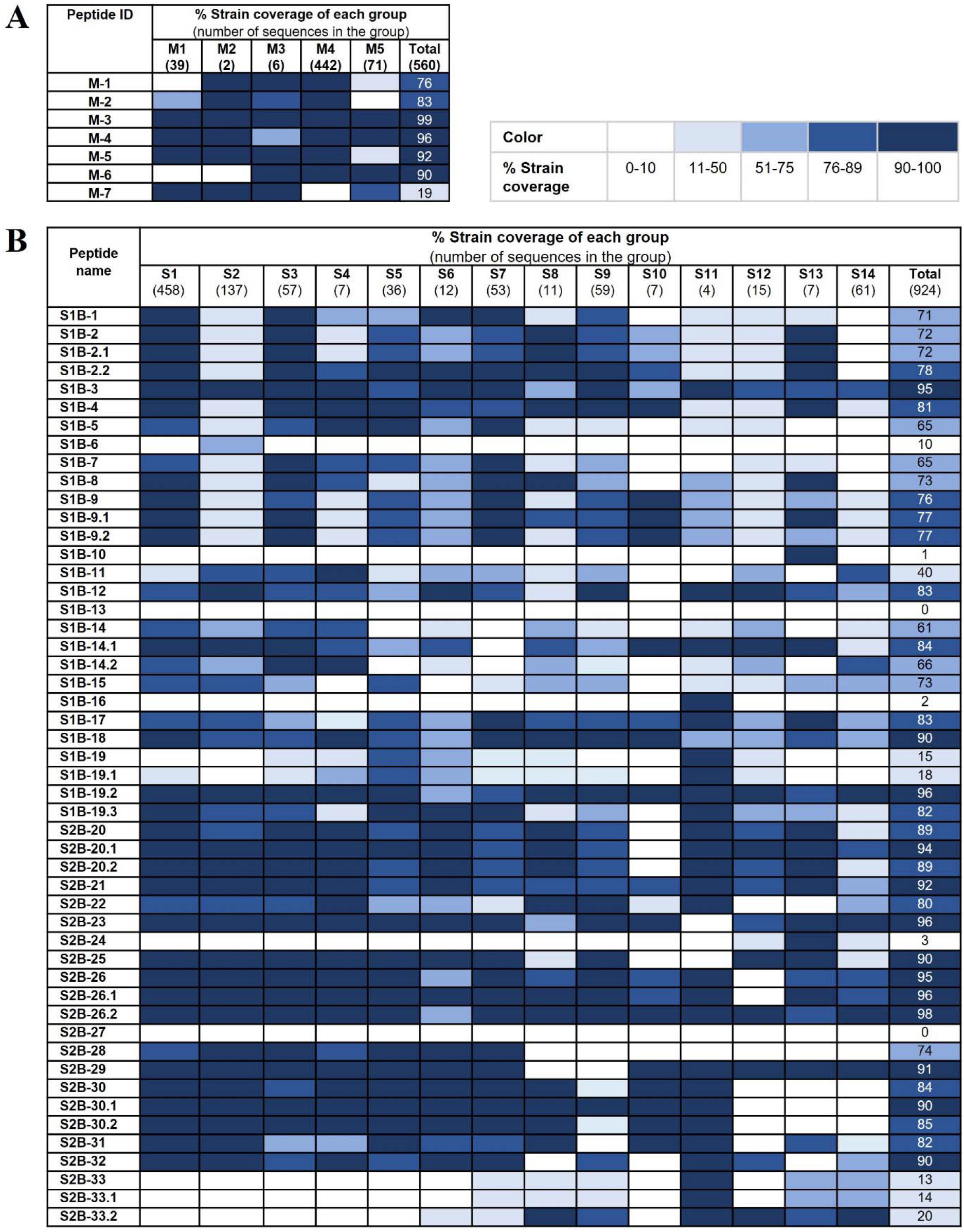
Conservancy of each predicted epitope among global PEDV strains. **(A** and **B)** Strain coverage of the predicted epitope of the M and S proteins, respectively. Percentage of strain coverage was calculated based on 100% sequence identity between the predicted epitope and the target sequence.

### B cell epitopes on the S2 subunit and M protein are more conserved than those on the S1 subunit

We further analyzed the strain coverage of each predicted epitope, as information towards epitope conservancy could be beneficial to the design and development of a vaccine, particularly a universal vaccine. For the M protein, 6 out of 7 predicted epitopes cover higher than 70% of 560 accession numbers with 4 epitopes (M-3, M-4, M-5 and M-6) having strain coverage greater than 90% (**Fig. 5A**). Analyzed with 924 accession numbers of the PEDV S protein, 34 out of 50 predicted epitopes of the S protein exhibited strain coverage greater than 70% (**Fig. 5B**). Among these 34 epitopes, 24 showed strain coverage higher than 80%, of which 8 and 16 are located on the S1 and S2 subunits, respectively, suggesting that the S1 subunit is more variable than the S2 subunit.

### Predicted B cell epitopes are recognized by PEDV-positive sera

To further validate the predicted linear B cell epitopes, ELISA was performed. PEDV-positive sera were prepared from F1 and F3 pigs raised in the farms and naturally infected with PEDV. Neutralizing activity of the pig sera were investigated using neutralization assay and only the serum samples with neutralizing antibody titers at least 32 (reciprocal serum dilution) were further used in ELISA analysis. A total of 12 F1 pigs and 11 F3 pigs showed neutralizing activity that passes the criteria (**Fig. 6**) and were subject to ELISA. Control sera were prepared from 10 uninfected F3 pigs raised in the animal research facility and their neutralizing antibody titer was confirmed to be lower than 32 by neutralization assay (**Supplementary Table 1**).

**Figure 6.**
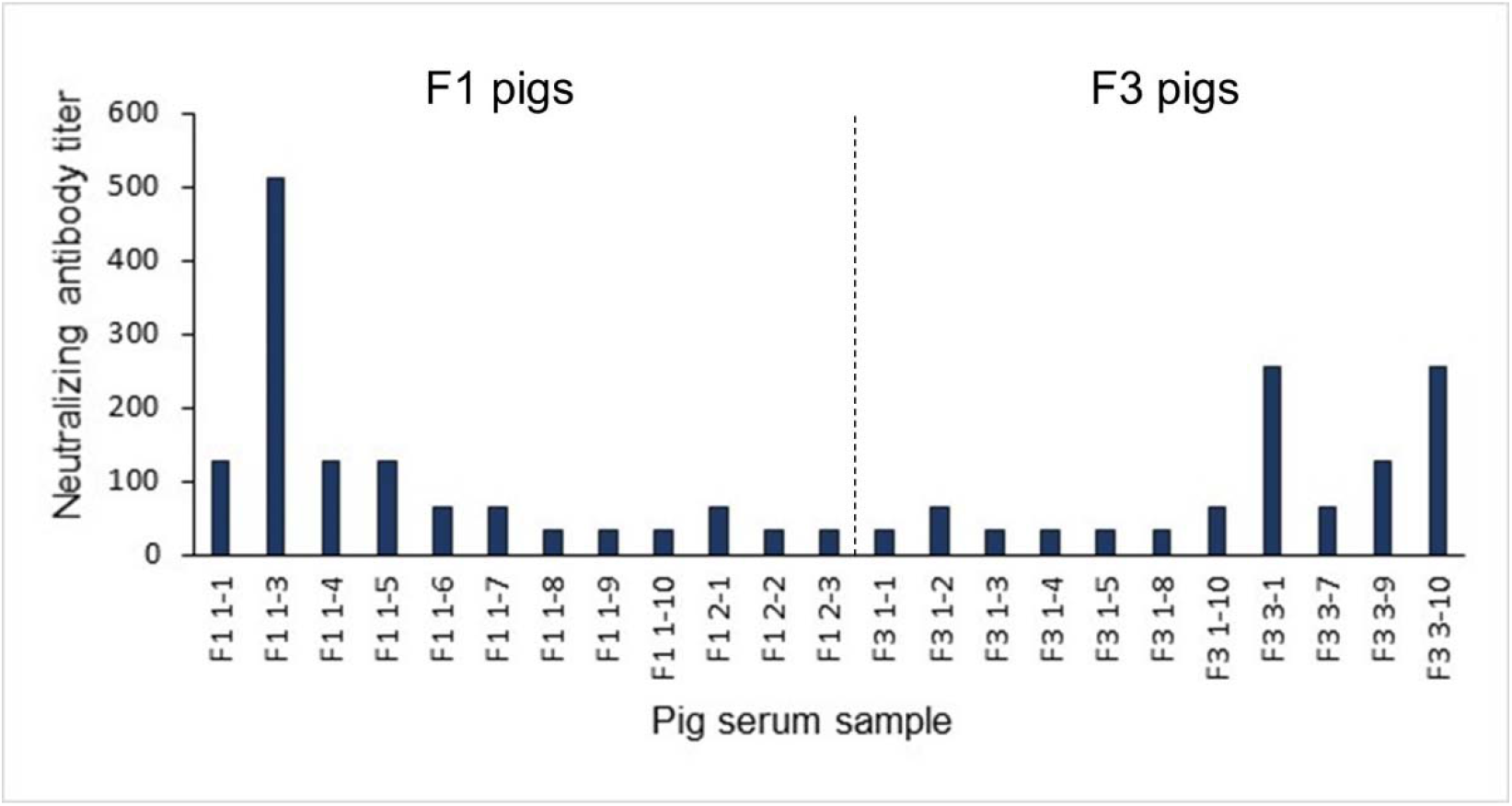
Neutralization titer of the pig sera. Pig sera were 2-fold serially diluted and incubated with PEDV (10 TCID_50_) before adding to Vero cells. PEDV infection was investigated by immunofluorescence staining using anti-PEDV monoclonal antibody at 3 days post-inoculation.

Three well known epitopes, SS2, SS6 and 2C10 (21, 24), were included in ELISA as reference epitopes. Reactivity of the peptides with their target antibodies was determined based on statistical analysis comparing OD450 value of the F1 and F3 pig sera to that of the control group (*p <* 0.05). All 7 peptides from the M protein showed reactivity with sera from at least one pig strain (**Fig. 7A**). Out of 50, 48 peptides of the S protein showed reactivity with the sera from at least one strain pig strain (**Fig. 7C**). Based on the analyses to determine immunodominant and subdominant epitopes (described in materials and methods), the epitopes S1B-1, S1B-2.2, S1B-3, S1B-9.2, S1B-14.2, S1B-19.1, S1B-19.3, S2B-22, S2B-25, S2B-29, and S2B-30.2 on the S protein and the epitope M-2 on the M protein were categorized as immunodominant epitopes, while the epitopes S1B-2, S1B-5, S1B-9.1, S1B-15, S1B-16, S1B-19.2, S2B-21, S2B-24, S2B-26.1 and S2B-32 were categorized as subdominant epitopes (**Fig. 7A-D, and Table 2)**. Two subdominant epitopes, S1B-15 and S1B-16, are located on the COE domain.

**Table 2.**
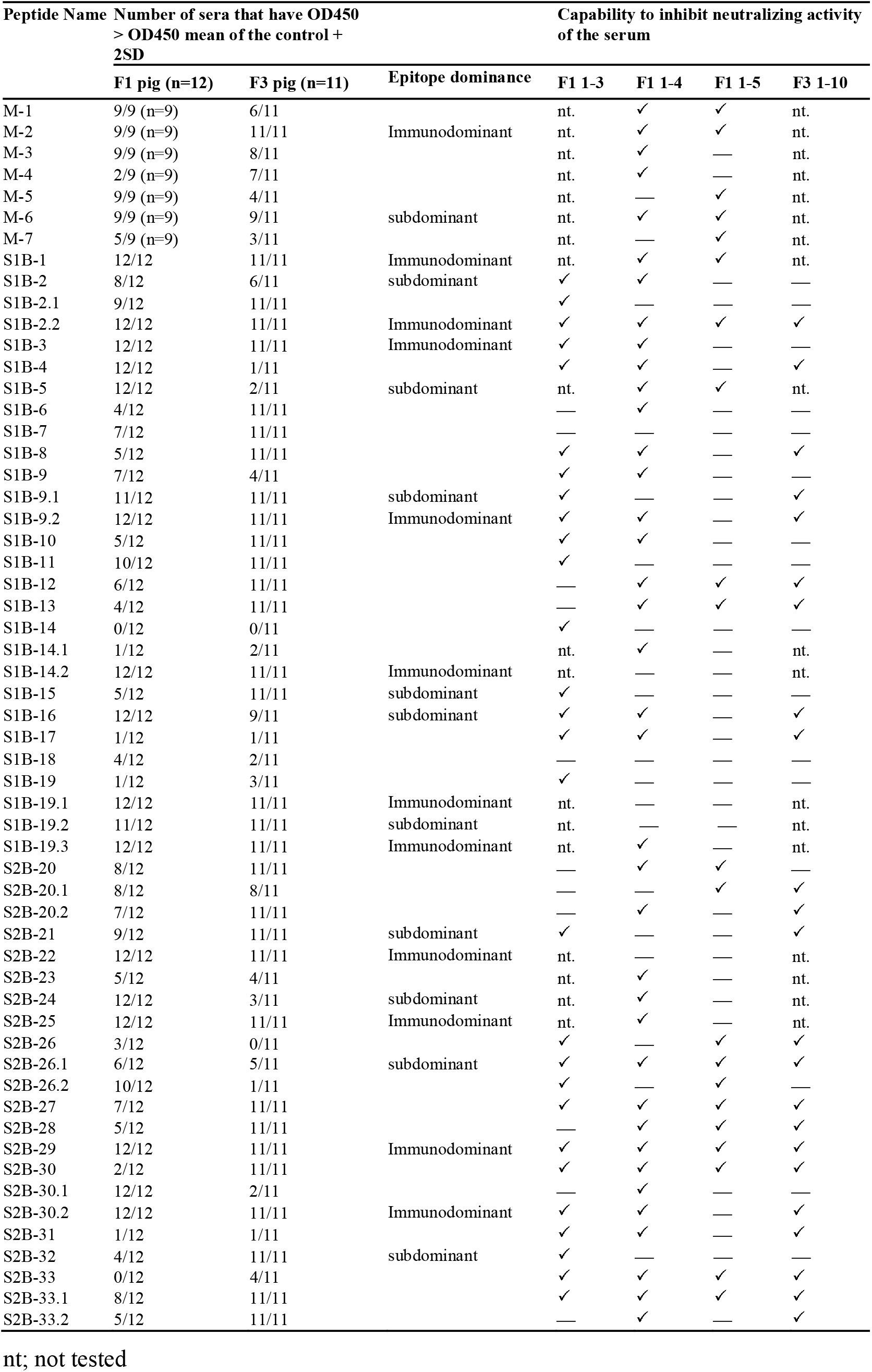
Potential as a neutralizing epitope determined by neutralizion-inhibition assay.

**Figure 7.**
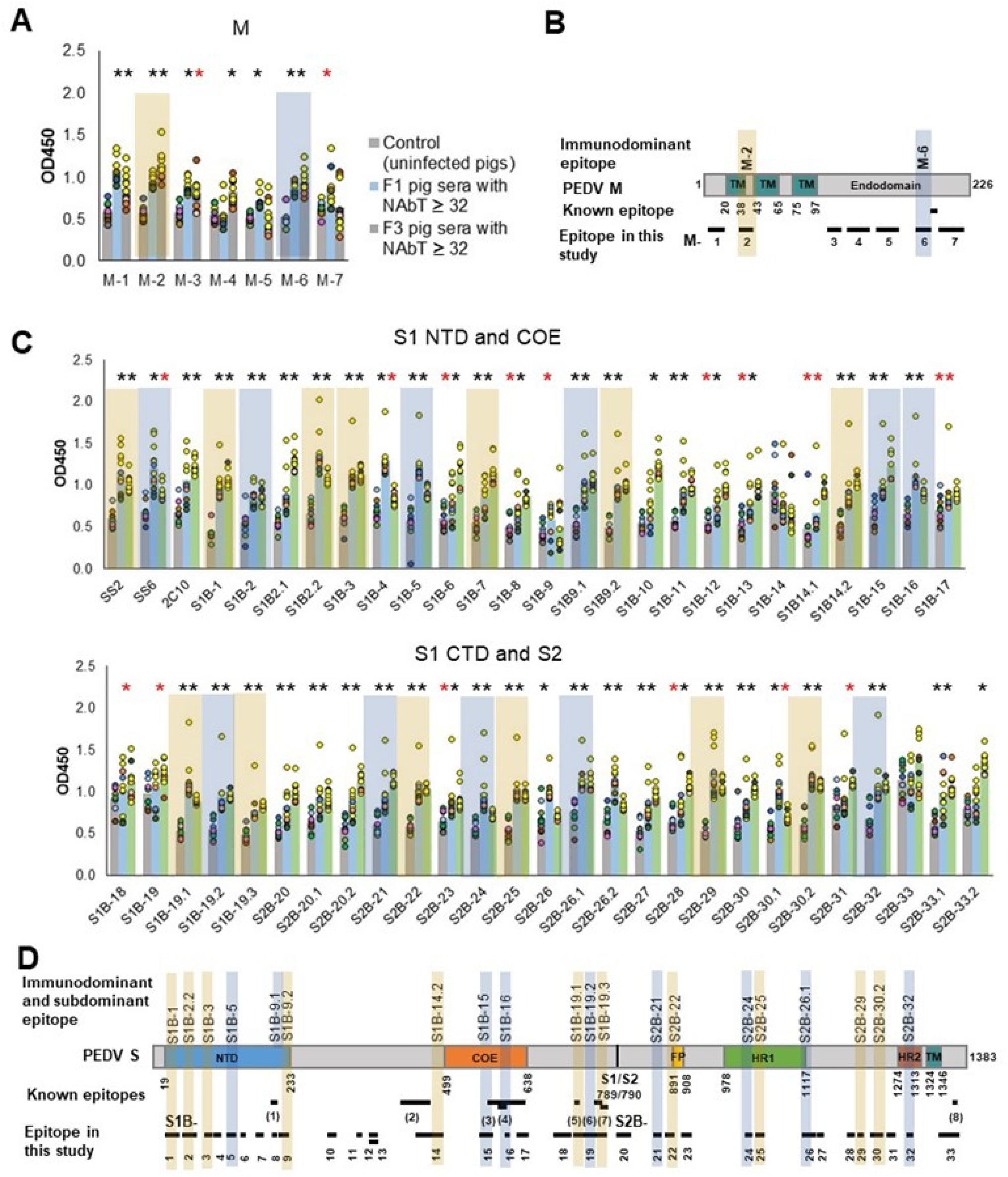
Antibody responses against predicted epitopes and landscapes of the immunodominant B cell epitopes. **(A** and **C**) Antibody responses against predicted B cell epitopes of the M and S proteins, respectively. ELISA was performed with PEDV-positive F1 (12 samples) and F3 (11 samples) pig sera whose neutralizing activity against PEDV had been confirmed. Sera from uninfected F3 pigs were used as a control. ELISA plates were coated with synthetic peptides corresponding to predicted epitopes as indicated in the X axis. Serum was diluted 1:50 and used in ELISA. The optical density at the wavelength of 450 nm was measured. The responses of serum samples with neutralizing antibody titer equal or higher than 64 are shown in yellow circle. Non-parametric Mann-Whitney U test was used in statistical analysis. * (red asterisk) indicates *p <* 0.05; * (black asterisk) indicates *p <* 0.001. Immunodominant and subdominant epitopes are indicated in yellow and blue box, respectively. **(B** and **D)** Landscape of the immunodominant B cell epitopes on the PEDV M and S proteins, respectively. Positions of amino acid residues in the M and S proteins are based on the sequence of PEDV CV777 (accession no. AAK38659.1). The N-terminal domain (NTD), fusion peptide (FP), heptad repeat regions (HR1 and HR2), and transmembrane domain (TM) are indicated in the diagram. All Immunodominant epitopes and some subdominant epitopes with their locations are indicated on the protein.

However, some of the B cell epitopes we identified here have already been reported. The S1B-19 epitope (aa. 719-772) is a part of the S1D domain (aa. 636-789) containing neutralizing epitopes (32). The S1B19.1 epitope (aa. 719-730) overlaps with the known epitope, S1D SE16 (33), while the S1B-19.3 epitope (aa. 748-772) consists of 2 known epitopes, SS2 (21) and SS6 (21). The S1B-8 epitope (aa. 203-211) corresponds to the known epitope, Peptide M (22), while the epitopes S2B-33 (aa. 1345-1379) contains the 2C10 epitope (34). The epitope S1B-14.1 (aa. 457-477) entirely overlaps with the neutralizing epitope (aa. 432-481) (35). In the COE region, all three epitopes (S1B-15 to S1B-17) are parts of the conformational epitope (572-<u>TLQSVND</u>YLSFSKFCVSTSLLASACTIDLF<u>GYPEFGSG</u>VKFTSLYFQFTK<u>GELITGTPKPLEGVT</u>-636) (35) as underlined. Additionally, the S1B-16 epitope also has 3 amino acids (GYP) overlapping with the C2-1 epitope (TSLLASACTIDLF<u>GYP</u>) (36). Besides these known epitopes, other B cell epitopes identified in our study are considered novel. Thus, epitopes S1B-1, S1B-2.2, S1B-3, S1B-9.2, S1B-14.2, S2B-22, S2B-25, S2B-29 and S2B-30.2 can be recognized as novel immunodominant epitopes. For the M protein, one B cell epitope named M14 (8), has been reported; however, this epitope does not overlap with the epitopes identified in our study. Therefore, all 7 B cell epitopes in the M protein we identified are novel epitopes.

### Novel neutralizing epitopes are verified by neutralization-inhibition assay

Neutralization ability of the antibodies against each epitope were further tested using neutralization-inhibition assay as shown in schematic representation in **Fig. 8A**. Four sera including F1 1-3, F1 1-4, F1 1-5 and F3 1-10 were used in the assay. Peptides corresponding to the known epitopes SS2, SS6 and 2C10 were used as references, and irrelevant peptides N(MHC) and E(CTL) were included as negative controls. All 3 reference peptides exhibited their ability to inhibit neutralizing activity of the sera, while irrelevant peptides did not (**Fig. 8B**). Notably, peptides from the M protein were tested with only 2 sera, F1 1-4, F1 1-5. While the F1 1-4 serum lost its neutralizing activity when incubated with peptides M-1, M-2, M-3, M-4 and M-6, neutralizing activity of serum F1 1-5 was depleted in the presence of peptides M-1, M-2, M-,5 M-6 and M-7 (**Fig. 8B** and **Supplementary Fig. 8-9)**. A combined result from both sera suggested that all 7 epitopes of the M protein had potential as neutralizing epitopes.

**Figure 8.**
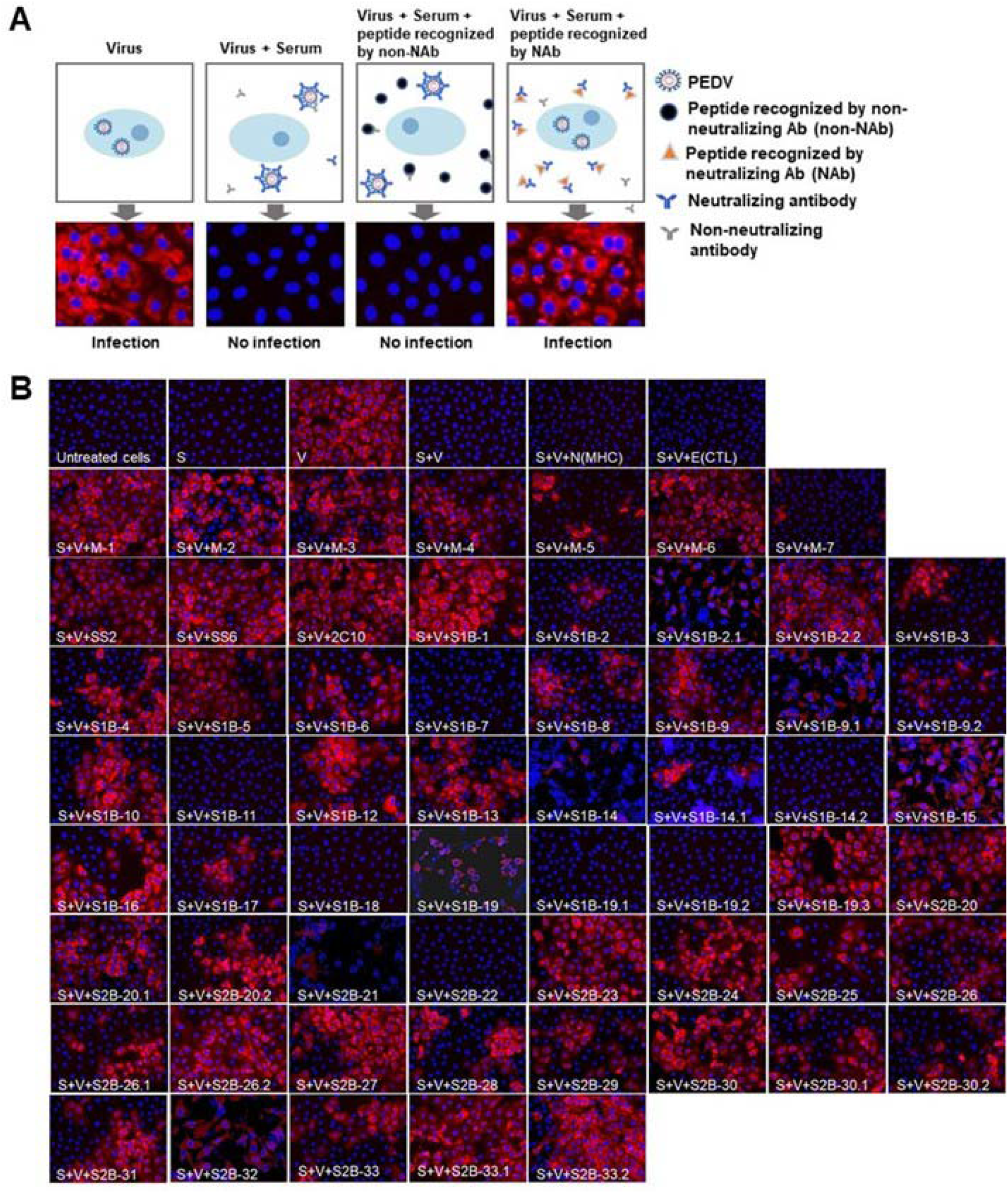
Neutralization-inhibition assay. **(A)** Schematic representation of neutralization-inhibition assay. **(B)** Inhibition of virus neutralization by synthetic peptides. Peptides (as indicated in the picture) were incubated with sera from PEDV-infected F1 and F3 pigs, prior to a further incubation with PEDV. The mixture was inoculated into Vero cells and incubated for 3 or 5 days. PEDV was detected using anti-PEDV monoclonal antibody and shown by red fluorescence. DAPI was used to determine nucleus (blue fluorescence). Immunofluorescence images were observed and taken under inverted fluorescence microscope. S: serum; V: virus; N(MHC) and E(CTL): irrelevant peptides.

In the presence of the S peptides, four pig sera showed different patterns of neutralization inhibition as summarized in **Table 2** and **Supplementary Fig. 7-10**. In **Fig. 8B**, neutralizing epitopes were concluded based on the combined results of all 4 sera. The peptide that inhibited neutralizing activity of at least one serum was considered a neutralizing epitope. Almost all peptides, except S1B-7, S1B-11, S1B-14.2, S1B-19.1, S1B-19.2 and S2B-22, could inhibit neutralizing activity of at least one serum. Hence, the epitopes S1B-1, S1B-2.2, S1B-3, S1B-9.2, S1B-19.3, S2B-22, S2B-25, S2B-29 and S2B-30.2 posed as both neutralizing epitopes as well as immunodominant epitopes. Importantly, antibodies recognizing the 3 epitopes in the COE domain, which are S1B-15, S1B-16 and S1B-17, were demonstrated with neutralization ability.

### Depiction of the identified B cell epitopes in the 3-D structure of the PEDV S protein

Surface representation of all epitopes is depicted on the prefusion structure of the trimeric S protein of the PEDV strain USA/Colorado/2013 (PDB: 6VV5) (37) using PyMOL 2.3.4. As shown in **Fig. 9A**, most of the epitopes are exposed on the surface, one main feature associated with B cell epitopes. Although coil is generally recognized as one main characteristic of linear B cell epitopes, B cell epitopes identified in our study were found to be in various structures including coil, alpha helix and beta sheet (**Fig. 9B**). Close-ups of the immunodominant epitopes and neutralizing epitopes in the COE revealed that these epitopes are either partly or entirely composed of coil structure and some epitopes also consist of other structures either alpha helix or beta sheet (**Fig. 9C-D**).

**Figure 9.**
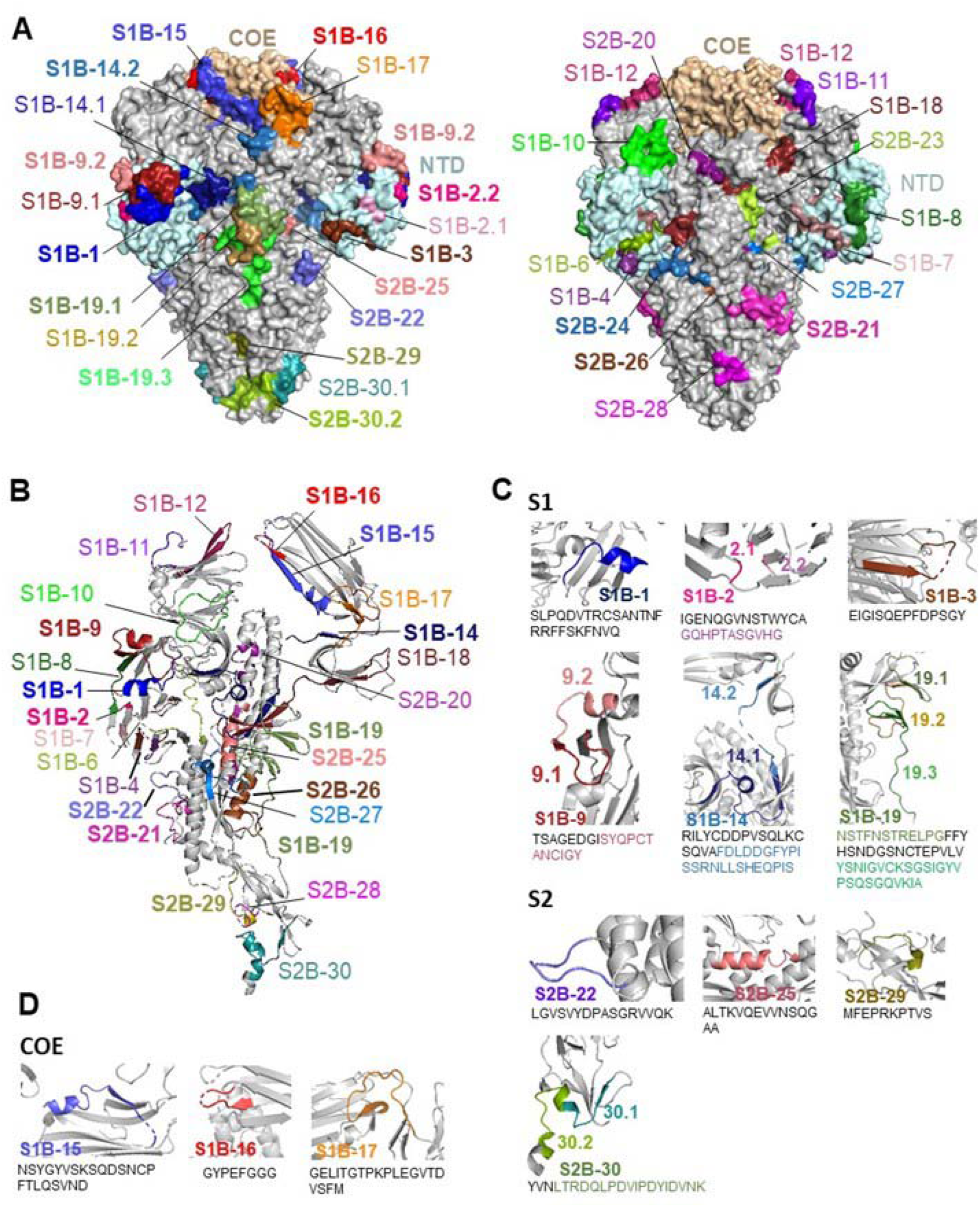
Surface representation and locations of the B cell epitopes on the S protein. **(A)** Surface representation of the B cell epitopes on the trimeric S protein. Left panel demonstrates immunodominant epitopes; right panel demonstartes subdominant epitopes. (**B**) Localization and structure of the epitopes on the monomeric S protein. **(C)** Close-ups of immunodominant epitopes. **(D)** Close-ups of neutralizing epitopes in the COE domain.

## DISCUSSION

Genetic diversity of PEDV, particularly in the S protein (6), has limited protective potency of the PEDV vaccine. Identification of neutralizing B cell epitopes that are conserved across all diverse PEDV strains will contribute to the development of a more effective universal vaccine to combat all diverse strains of PEDV. So far, 8 B cell epitopes have been identified from 5 regions of the PEDV S protein. With the aim to identify B cell epitopes from diverse strains of PEDV, immunoinformatics approach allows us to primarily screen for candidate B cell epitopes from a large number of sequences retrieved from the database. By using multiple prediction tools, none of these prediction tools gave the prediction result that covers all 4 well known epitopes (peptide M (22), SS2 (21), SS6 (21) and 2C10 (34)) when they were used alone. Our work is in agreement with the previous study showing that prediction using 2 or more tools increased both accuracy and probability in defining a genuine B cell epitope (38). Importantly, immunoinformatics methods we developed in this study allowed us to identify candidate B cell epitopes from all functionally important regions, which include all reported epitopes of the PEDV S protein. ELISA test confirmed reactivity of the predicted peptides with PEDV-positive sera. However, the patterns of antibody responses in 2 different pig strains, F1 and F3, were different with some peptides. This may be influenced by several factors such as pig genetic background, age, duration and dose of infection. Moreover, it is also possible that the infection in each farm are caused by different strains of PEDV, as PEDV found in different regions of Thailand has been reported with genetic diversity (39).

Statistical analysis of antibody responses in both F1 and F3 pig sera defined 11 immunodominant within the PEDV S proteins. Interestingly, although PEDV (alphacoronavirus) and SASR-CoV-2 (betacoronavirus) belong to different genus, the landscape of immunodominant epitopes on the S protein of these two coronaviruses remarkably resembles. Epitopes (i) S1B-1, (ii) S1B-19 (19.1 and 19.3), (iii) S2B-22, and (iv) S2B-29 and S2B-30.2 are located at (i) the N terminus next to the signal sequence, (ii) CTD, upstream region of the S1/S2 cleavage site, (iii) FP region, and (iv) upstream region of the HR2, respectively, which have also been indicated to be the main sites of immunodominant epitopes on the SARS-CoV-2 S protein studied by our group (manuscript submitted). and Li *et.al*. (2021) (40). Moreover, the epitopes S1B-15, S1B-16 and S1B-17, which are located within the COE domain, also mirror the 3 most dominant epitopes within the S RBD of SARS-CoV-2 identified in our previous work (manuscript submitted). Additionally, the C-terminal endodomain of the S protein is another main site accommodating the immunodominant B cell epitope of SARS-CoV-2 (40–42) as well as the neutralizing epitope 2C10 with the GPRLQPY motif (epitope S2B-33 in this study) of PEDV (34).

Neutralizing activity of the antibodies targeting these novel epitopes was addressed using neutralization-inhibition assay, which has been previously used to identify neutralizing epitopes from SARS-CoV-2 (41) and coxsackievirus A16, a causative agent of human, foot, hand and mouth disease (43). However, we observed that inhibitory effect of peptide on different sera were different for some peptides. This may be because the levels of antibodies against each epitope vary in different sera. Surprisingly, we did not see inhibitory effect of 3 immunodominant epitopes, S1B-14.2, S1B-19.1, and S2B-22, on any sera. This result can be explained by 2 possibilities: (i) these epitopes are indeed non-neutralizing epitopes recognized by non-neutralizing antibodies, and (ii) the peptides cannot compete with the S protein present on the virus particle in binding to their cognate antibodies; thus, neutralizing activity of the serum is maintained.

Neutralizing epitopes we identified are located at the regions that are functionally important during virus infection. Epitopes S1B-1 to S1B-9 are located at the N-terminal domain (NTD) of the S1 subunit, the region that involves in binding to sialoglycoconjugates receptor on the host cell (44, 45). The neutralizing epitope S1B-10 are located at the S1 region that plays an important role in PEDV attachment to host cell using sugar-binding activity (46). Moreover, the neutralizing epitope S1B 14.1 overlaps with the C-terminal sequence of the conformational epitope (aa. 435-485) previously identified in the PEDV PT strain (35), suggesting that the overlapping sequence (457-RILYCDDPVSQLKCSQVAFDL-477) may function as a core neutralizing epitope in that region. Within COE domain, 3 neutralizing epitopes we identified, S1B-15, S1B-16 and S1B-17, may be the main targets for neutralizing antibody recognition in the conformational and neutralizing epitope (aa. 572-636) previously identified (35). The S1D domain located in the S1 CTD, which also includes our neutralizing epitope S1B-19, is well documented as an immunodominant epitope/domain and a target of neutralizing antibodies (10, 21, 47).

While most of the known neutralizing B cell epitopes are located in the S1 subunit, information of B cell epitopes in the S2 subunit are limited with only one neutralizing epitope identified. The 2C10 epitope located at the C-terminus of the S2 domain has been identified as neutralizing epitope (23, 24). In this study, in addition to the 2C10 epitope, more neutralizing B cell epitopes in the S2 subunit were identified. The neutralizing epitope S2B-20 is located next to the S1/S2 cleavage site; thus, antibody binding to this region may interfere S1/S2 cleavage, resulting in a loss of viral infectivity. Coronavirus S2 subunit consists of Fusion peptide (FP) and Heptad repeat regions (HR) that play important roles in cell membrane fusion (48). In a betacoronavirus, SARS-CoV-2, the epitopes located in the FP and HR2 regions have been identified as immunodominant and neutralizing epitope (49). In our study, the neutralizing epitopes S2B-21 and S2B-23 are located in close proximity to the FP, a region responsible for protease cleavage and membrane fusion (45); thus, antibody binding to this region could affect these processes during viral cell entry. Neutralizing epitopes S2B-24, 25 and 26 are located within the HR1 region, whereas neutralizing epitopes S2B-27, 28, 29, 30 and 31 are located in the upstream region of the HR2 and S2B-32 is located within the HR2. The HR1 and HR2 domains of coronaviruses play important roles in membrane fusion (45); thus, blocking these regions with antibodies may result in inhibition of the membrane fusion process.

As our B cell epitopes are predicted based on consensus sequence of each group of the S and M proteins, the epitope sequence may not 100% match with all PEDV strains/variants. However, some epitopes are highly conserved as evidenced by high percentage of strain coverage among global PEDV strains. Compared to the epitopes in the S protein, epitopes in the M protein were found to be more conserved. Within the S protein, epitopes in the S2 subunit are more conserved than those in the S1 subunit. These conserved epitopes may serve as candidates for development of a multi-epitope universal vaccine.

Taken together, the method combining immunoinformatics with immunoassays enabled identification of novel neutralizing linear B cell epitopes on the PEDV S and M proteins. Even though these B cell epitopes are derived from consensus sequence, some of which are highly conserved among the global PEDV strains, which represent a promising vaccine target for development of a universal epitope-based vaccine as well as for antibody detection. Importantly, the immunoinformatics method developed in this study can serve as a useful tool for prediction of linear B cell epitopes from any protein of interest.

## MATERIALS AND METHODS

### Protein sequence retrieval and phylogenetic analysis

Amino acid sequences of the PEDV S and M proteins were retrieved from National Center for Biotechnology Information (NCBI) (https://www.ncbi.nlm.nih.gov/protein/). Retrieved protein sequences were analyzed individually based on a defined name and length using Sublime Text3 program. Based on sequence similarity, the amino acid sequences of the S and M protein were grouped using alignment and phylogenetic tree tools in SeaView program (version 4). In SeaView program, these amino acid sequences were aligned using MUSCLE and grouped using tree based on distance analysis, NJ method, ignore gap and bootstrap 1000 replication. Consensus sequence of each group was generated using MUSCLE method in Unipro UGENE program (version 1.24.2; September 1, 2016) (50, 51). These consensus sequences were then subjected to epitope prediction.

### B cell epitope prediction

Immunoinformatics tools were exploited to predict B cell epitopes and properties of the amino acid residues. Firstly, linear B cell epitope prediction was performed using BepiPred-2.0 (http://www.cbs.dtu.dk/services/BepiPred/), which predicts linear B cell epitopes from a protein sequence, using a Random Forest algorithm trained on epitopes and non-epitope amino acids determined from crystal structures, followed by performing sequential prediction smoothing (52). In our study, the residues with the threshold score of epitope probability above 0.5 were considered as parts of a B cell epitope. Putative B cell epitopes were selected based on the region with at least 6 consecutive residues predicted to have epitope probability above 0.5. Notably, BepiPred-2.0 also predicts and provides accessibility and coil probability of each amino acid residue. In addition, other properties of the proteins were also characterized using multiple immunoinformatics tools provided by IEDB (http://tools.iedb.org/bcell/), except IUPred. IUPred (https://iupred.elte.hu/) was used to predict intrinsically unstructured proteins which infers to coil probability (53). The method of Emini (54) was used to predict surface accessibility, while the method of Kolaskar & Tongaonkar, semi-empirical method (55), was used to predict antigenicity determinant. Hydrophilicity of the protein was predicted using the method of Parker. The results of prediction from each method were aligned and 3 methods for identifying and selecting candidate B cell epitopes were created based on the 4 known B cell epitopes derived from PEDV protein as described in the result part.

### Epitope conservancy analysis

As epitopes were predicted based on consensus sequences of each group of the PEDV S (14 groups) and M (5 groups) proteins. Epitope conservancy was analyzed by calculating percent strain coverage of the predicted epitopes in comparison to all of the sequences in the group using the formula below.

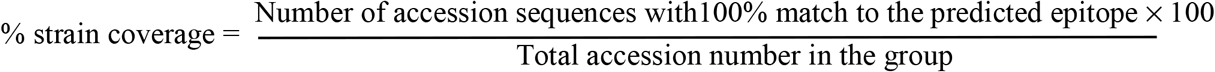

### Peptide synthesis and preparation

Peptides corresponding to immunoassay were chemically synthesized (Mimotopes). All synthetic peptides were (i) 57 predicted epitopes, (ii) 3 known reference B cell epitopes, SS2 (YSNIGVCK) (21), SS6 (LQDGQVKI) (21), and 2C10 (GPRLQPY) (24) and (iii) an irrelevant peptide, N(MHC) (LADSYEITY) and E(CTL) (AVYTPIGRLY). Synthetic peptides were dissolved to the concentration of 3 nmol/μl (3 mM) as peptide stock in sterile distilled water containing 0.1% acetic acid. Peptide stock were aliquoted and stored at −20 °C until used.

### Animal ethics and preparation of pig sera

All procedures involving collecting samples from pigs raised in the farms were performed in accordance with guidelines of Institutional Animal Care and Use Committee (IACUC) of Institute for Animals for Scientific Purpose Development and carried out under the regulation and permission of King Mongkut’s University of Technology Thonburi IACUC protocol No. KMUTT-IACUC-2019/018. For the control group, pigs were raised in Mahidol University’s animal research facility. All procedures were carried out under the regulation and permission of Faculty of Veterinary Science, Mahidol University IACUC protocol No. MUVS-KA-2021-01-01.

Blood samples (10 ml) were collected from pigs naturally infected with PEDV. Infected pigs were observed and diagnosed by the veterinarian based on signs and symptoms of PED. Pigs showing PED signs or had been diagnosed with PED within the past 4 weeks were used as subjects. Bloods were collected from 2 different pig strains: F1 sow (Danish Landrace, age 35-40 weeks old) and F3 fattening pigs (Danish Landrace x Large white x Danish Duroc, age 7-12 weeks old). Blood were collected from 13 female F1 pigs and 14 either male or female pigs raised in 3 different farms. Control sera were prepared from 10 uninfected F3 pigs (age 7 weeks old) raised in Mahidol University’s animal research facility. Bloods were left to clot at room temperature for approximately 2 h, followed by centrifugation at 1,500 x g for 10 min in a refrigerated centrifuge to separate the serum. Serum was aliquoted and separated to a small portion in new tubes and stored at −20 °C until used.

### Cell and PEDV preparation

PEDV used in this study is a lineage of PEDV CV777. Vero cells were cultured in complete DMEM medium supplemented with 10% fetal bovine serum (FBS, HyClone) and 1% penicillin/streptomycin (Pen/Strep, gibco), and then incubated at 37 °C with 5% CO_2_. The seed virus was sequentially propagated in Vero cell with PEDV medium (DMEM supplemented with 0.02% yeast extract (HIMEDIA), 10 μg/ml of trypsin (SIGMA), and 0.3% tryptose phosphate broth (tryptose, 20 g/L; dextrose, 2 g/L; sodium chloride, 5 g/L; disodium hydrogen phosphate, 2.5 g/L) (35). To prepare PEDV stock, Vero cells were cultured in 10 flasks of T175. When the cell confluency was over 90%, old medium was removed and the cells were infected with PEDV resuspended in 20 ml of PEDV medium. When cytopathic effect (CPE) reached 70%, medium was harvested and pooled. Sodium chloride (NaCl) was added to the medium at a final concentration of 0.5 M and polyethylene glycol (PEG) was then added to a final concentration of 8%. After incubated overnight, PEDV was harvested by centrifugation at 8,000 rpm, 20 min. The viral pellet was next resuspended in 5 ml of endotoxin-free PBS, aliquoted in a small volume to new tubes and stored at −80 °C. Next, virus titer was determined using 50% tissue culture infective dose (TCID_50_) method. Virus suspension was diluted in ten-fold serial dilutions and pipetted into ten wells of confluent Vero cells. After 3 days post infection, PEDV-infected cells were observed and counted using immunofluorescence staining assay. TCID_50_ was next calculated.

### Immunofluorescence staining assay

Three days post-transfection, Vero cells in 96-well plate were fixed with 1% formaldehyde at RT for 15 min and then washed with PBS once. Cells were permeabilized with cold 90% methanol and then incubated at 4 °C for 5 min. Cells were washed once with PBS and blocked with 2% FBS/PBS for 1 h at RT. The blocking solution in each well was replaced with 50 μl of mouse anti-PEDV (Median Diagnostic) diluted to 1:1,000 in PBS. Following an incubation for 2 h at RT, the cells were washed twice with PBS. Secondary antibody, donkey anti-mouse IgG conjugated with Alexa Fluor^®^ 594 (Abcam) diluted 1:1,000 in PBS, was added to the cell and incubated for 1 h at RT. The cells were washed twice with PBS and then stained the nucleus with DAPI (Sigma) diluted 1:1,000 in PBS and incubated at RT for 30 min, followed by washing once with PBS. Fluorescence images were directly examined under an inverted fluorescence microscope (Olympus DP74).

### ELISA

Synthetic peptides (0.75 nmole in 50 μl PBS/per well) were added into 96-well ELISA microplates (Greiner bio-one). After an overnight incubation at 4 °C, plates were washed 3 times using PBS containing 0.05% Tween 20 (PBST). Plates were blocked using 100 μl of PBST containing 5% FBS and incubated with agitation at room temperature (RT) for 1 h. Pig sera diluted 1:50 in PBST containing 1% FBS were added to the plates. After a 2-h incubation at RT, the plates were washed 3 times with PBST and goat anti-pig IgG HRP antibody (Abcam) diluted 1:10,000 in PBST containing 1% FBS was added to the well (60 μl/well), followed by an incubation at RT for 90 minutes. After a 3-time wash with PBST, TMB substrate (BioLegend) was added (70 μl/well) and the plate was incubated in dark at RT for 30 min, allowing the color to develop. The reaction was stopped by adding 35 μl of 1 M sulfuric acid (H_2_SO_4_) and the optical density was measured at a wavelength of 450 (OD450) (MULTISKAN FC, Thermo scientific).

### Antibody neutralization assays

All pig serum samples were 2-fold serially diluted with DMEM containing 1% Pen/Strep to the dilution ranging from 1:32 to 1:4,096. PEDV at the concentration of 200 TCID_50_/ml was prepared using PEDV medium. PEDV (10 TCID_50_ in 50 μl) were mixed with 50 μl of diluted serum or medium (control) and incubated at 37 °C with 5% CO_2._ After, a 1-h incubation, the mixture of 100 μl was transferred into 96-well plates containing approximately 90% confluent monolayer of Vero cells. Each condition was performed in duplicate. After a 3-h incubation, the medium was removed and replaced with new PEDV medium. At day 3 post-infection, immunofluorescence staining assay was performed and the number of infected cells were counted. End point neutralizing antibody titer was determined based on the serum dilution that completely inhibited virus infection.

### Neutralization-inhibition assay

Neutralization-inhibition assay was conducted as previously described by Shi et al. (2013) (43) with some modifications. Sera from 3 F1 pigs (F1 1-3, F-1 1-4 and F1 1-5) and one F3 pig (F3 1-10) were subject to the assay. Serum was 2-fold serially diluted with DMEM containing 1% antibiotic to the dilution ranging from 1:32 to 1:256. PEDV was diluted in PEDV medium to 2,000 TCID_50_/ml. Synthetic peptides were mixed with the diluted serum to a final concentration of 200 nmol/ml in a volume of 50 μl. After an incubation at 37 °C for 1 h, PEDV (100 TCID_50_ in 50 μl) was added to the peptide-serum mixture. After an incubation at 37 °C for 1 h, the mixture (100 μl) was transferred to Vero cells grown in 96-well plates. Following an incubation at 37 °C with 5% CO_2._for 3 h, the virus suspension was removed from the well and new medium was added into the wells. In addition, the conditions that the Vero cells were incubated with (i) medium alone, (ii) serum alone, (iii) PEDV alone, and (iv) the mixture of PEDV and serum, were also included in the study as control conditions. All conditions were performed in duplicate. Following a 3-day (F-1 1-4 and F1 1-5) or 5-day (F1 1-3 and F3 1-10) incubation, PEDV infection in Vero cell was investigated using immunofluorescence staining assay. Neutralization inhibition mediated by a peptide was determined at the endpoint neutralizing antibody titer.

### Labelling of neutralizing B cell epitopes

Putative neutralizing B cell epitopes were aligned and compared with full-length S protein of PEDV USA/Colorado/2013 strain for identification of epitope position. To localize each epitope within trimeric PEDV S protein, prefusion structure of PEDV spike named 6VV5 (37) was used as a model. All predicted epitopes were depicted PyMOL 2.3.4 program.

### Statistical analysis

The statistical significance of the samples in different groups was analyzed using SPSS 22 for Windows software (SPSS, USA). Data analyses were performed using non-parametric Mann-Whitney U test. The difference between the 2 groups was determined based on *p <* 0.05. Immunodominant and subdominant epitopes were defined with following criteria. In response to a particular peptide, if all sera from both F1 and F3 pigs exhibited OD450 value higher than OD450 mean of the control group + 2SD, such peptide was considered an immunodominant. On the other hand, if all sera from both F1 and F3 pigs exhibited OD450 value higher than OD450 mean of the control group + 1SD, such peptide was considered a subdominant epitope.

## Data Availability

Additional data in this study are provided in the Supplementary information. Additional supporting data and materials are available from the corresponding author Y.M.R. on request.

## Authors’ contributions

K.P.: designed and performed experiments, analyzed and interpreted data, wrote an original draft and revised the manuscript. M.R.: supervised immunoinformatics methods and experimental design. P.M.: provided materials and reagent. K.K.: prepared uninfected pig sera. T.H.: provided financial support, materials and reagents. P.J.: collected blood samples from PEDV-infected pigs. Y.M.R.: designed experiments, analyzed and interpreted data, supervised to K.P., reviewed and edited the manuscript. All authors read manuscript and gave comments.

## Acknowledgements

We would like to thank B.F. Feed Company Limited for providing financial support and providing blood samples from PEDV-infected pigs. Kanokporn Polyiam’s PhD study is funded by Development and Promotion of Science and Technology Talents Project (DPST).

## References

1. Paarlberg PL. Updated estimated economic welfare impacts of porcine epidemic diarrhea virus (PEDV). Purdue University, Department of Agricultural Economics, Working Papers. 2014;14(4):1–38.

2. Song D, Park B. Porcine epidemic diarrhoea virus: a comprehensive review of molecular epidemiology, diagnosis, and vaccines. Virus genes. 2012;44(2):167–75.

3. Huang Y-W, Dickerman AW, Piñeyro P, Li L, Fang L, Kiehne R, et al. Origin, evolution, and genotyping of emergent porcine epidemic diarrhea virus strains in the United States. MBio. 2013;4(5):e00737–13.

4. Sun RQ CR, Chen YQ, Liang PS, Chen DK, Song CX. Outbreak of porcine epidemic diarrhea in suckling piglets, China. Emerging infectious diseases. 2012;18(1):161–3.

5. Chaochao Lv, Yan Xiao, Xiangdong Li, Tian K. Porcine epidemic diarrhea virus: current insights. Virus adaptation and treatment. 2016;8:1–12.

6. Lee C. Porcine epidemic diarrhea virus: an emerging and re-emerging epizootic swine virus. Virology journal. 2015;12(1):193.

7. Pollard AJ, Bijker EM. A guide to vaccinology: from basic principles to new developments. Nat Rev Immunol. 2021;21(2):83–100.

8. Zhang Z, Chen J, Shi H, Chen X, Shi D, Feng L, et al. Identification of a conserved linear B-cell epitope in the M protein of porcine epidemic diarrhea virus. Virology journal. 2012;9(1):225.

9. Arndt AL, Larson BJ, Hogue BG. A conserved domain in the coronavirus membrane protein tail is important for virus assembly. J Virol. 2010;84(21):11418–28.

10. Okda FA, Lawson S, Singrey A, Nelson J, Hain KS, Joshi LR, et al. The S2 glycoprotein subunit of porcine epidemic diarrhea virus contains immunodominant neutralizing epitopes. Virology. 2017;509:185–94.

11. Chang CY, Cheng IC, Chang YC, Tsai PS, Lai SY, Huang YL, et al. Identification of Neutralizing Monoclonal Antibodies Targeting Novel Conformational Epitopes of the Porcine Epidemic Diarrhoea Virus Spike Protein. Sci Rep. 2019;9(1):2529.

12. Bosch BJ, van der Zee R, de Haan CA, Rottier PJ. The coronavirus spike protein is a class I virus fusion protein: structural and functional characterization of the fusion core complex. J Virol. 2003;77(16):8801–11.

13. Shirato K, Maejima M, Matsuyama S, Ujike M, Miyazaki A, Takeyama N, et al. Mutation in the cytoplasmic retrieval signal of porcine epidemic diarrhea virus spike (S) protein is responsible for enhanced fusion activity. Virus Res. 2011;161(2):188–93.

14. Chang S-H, Bae J-L, Kang T-J, Kim J, Chung G-H, Lim C-W, et al. Identification of the epitope region capable of inducing neutralizing antibodies against the porcine epidemic diarrhea virus. Molecules and cells. 2002;14(2):295–9.

15. Wicht O, Li W, Willems L, Meuleman TJ, Wubbolts RW, van Kuppeveld FJ, et al. Proteolytic activation of the porcine epidemic diarrhea coronavirus spike fusion protein by trypsin in cell culture. J Virol. 2014;88(14):7952–61.

16. Matsuyama S, Taguchi F. Two-step conformational changes in a coronavirus envelope glycoprotein mediated by receptor binding and proteolysis. J Virol. 2009;83(21):11133–41.

17. Sun YG, Li R, Xie S, Qiao S, Li Q, Chen XX, et al. Identification of a novel linear B-cell epitope within the collagenase equivalent domain of porcine epidemic diarrhea virus spike glycoprotein. Virus Res. 2019;266:34–42.

18. Do VT, Jang J, Park J, Dao HT, Kim K, Hahn TW. Recombinant adenovirus carrying a core neutralizing epitope of porcine epidemic diarrhea virus and heat-labile enterotoxin B of Escherichia coli as a mucosal vaccine. Arch Virol. 2020;165(3):609–18.

19. Li Q, Peng O, Wu T, Xu Z, Huang L, Zhang Y, et al. PED subunit vaccine based on COE domain replacement of flagellin domain D3 improved specific humoral and mucosal immunity in mice. Vaccine. 2018;36(11):1381–8.

20. Ma S, Wang L, Huang X, Wang X, Chen S, Shi W, et al. Oral recombinant Lactobacillus vaccine targeting the intestinal microfold cells and dendritic cells for delivering the core neutralizing epitope of porcine epidemic diarrhea virus. Microb Cell Fact. 2018;17(1):20.

21. Sun D, Feng L, Shi H, Chen J, Cui X, Chen H, et al. Identification of two novel B cell epitopes on porcine epidemic diarrhea virus spike protein. Vet Microbiol. 2008;131(1–2):73–81.

22. Cao L, Ge X, Gao Y, Zarlenga DS, Wang K, Li X, et al. Putative phage-display epitopes of the porcine epidemic diarrhea virus S1 protein and their anti-viral activity. Virus genes. 2015;51(2):217–24.

23. Cruz DJM, Kim C-J, Shin H-J. Phage-displayed peptides having antigenic similarities with porcine epidemic diarrhea virus (PEDV) neutralizing epitopes. Virology. 2006;354(1):28–34.

24. Cruz DJM, Kim C-J, Shin H-J. The GPRLQPY motif located at the carboxy-terminal of the spike protein induces antibodies that neutralize Porcine epidemic diarrhea virus. Virus research. 2008;132(1):192–6.

25. Dhanda SK, Usmani SS, Agrawal P, Nagpal G, Gautam A, Raghava GP. Novel in silico tools for designing peptide-based subunit vaccines and immunotherapeutics. Briefings in bioinformatics. 2017;18(3):467–78.

26. Shey RA, Ghogomu SM, Esoh KK, Nebangwa ND, Shintouo CM, Nongley NF, et al. In-silico design of a multi-epitope vaccine candidate against onchocerciasis and related filarial diseases. Scientific reports. 2019;9(1):4409.

27. Adam KM. Immunoinformatics approach for multi-epitope vaccine design against structural proteins and ORF1a polyprotein of severe acute respiratory syndrome coronavirus-2 (SARS-CoV-2). Trop Dis Travel Med Vaccines. 2021;7(1):22.

28. Behmard E, Soleymani B, Najafi A, Barzegari E. Immunoinformatic design of a COVID-19 subunit vaccine using entire structural immunogenic epitopes of SARS-CoV-2. Sci Rep. 2020;10(1):20864.

29. IEDB. Immune Epitope Database and Analysis Resource [cited 2017 February 10]. Available from: http://www.iedb.org/.

30. Kringelum JV, Nielsen M, Padkjær SB, Lund O. Structural analysis of B-cell epitopes in antibody: protein complexes. Molecular immunology. 2013;53(1):24–34.

31. Ponomarenko JV, Van Regenmortel MH. B cell epitope prediction. Structural bioinformatics. 2009;2:849–79.

32. Sun DB, Feng L, Shi HY, Chen JF, Liu SW, Chen HY, et al. Spike protein region (aa 636789) of porcine epidemic diarrhea virus is essential for induction of neutralizing antibodies. Acta Virol. 2007;51(3):149–56.

33. Kong N, Meng Q, Jiao Y, Wu Y, Zuo Y, Wang H, et al. Identification of a novel B-cell epitope in the spike protein of porcine epidemic diarrhea virus. Virology Journal. 2020;17(1):1–9.

34. Cruz DJ, Kim CJ, Shin HJ. Phage-displayed peptides having antigenic similarities with porcine epidemic diarrhea virus (PEDV) neutralizing epitopes. Virology. 2006;354(1):28–34.

35. Chang C-Y, Cheng I-C, Chang Y-C, Tsai P-S, Lai S-Y, Huang Y-L, et al. Identification of Neutralizing Monoclonal Antibodies Targeting Novel Conformational Epitopes of the Porcine Epidemic Diarrhoea Virus Spike Protein. Scientific reports. 2019;9(1):2529.

36. Sun Y-g, Li R, Xie S, Qiao S, Li Q, Chen X-x, et al. Identification of a novel linear B-cell epitope within the collagenase equivalent domain of porcine epidemic diarrhea virus spike glycoprotein. Virus research. 2019;266:34–42.

37. Kirchdoerfer RN, Bhandari M, Martini O, Sewall LM, Bangaru S, Yoon K-J, et al. Structure and immune recognition of the porcine epidemic diarrhea virus spike protein. Structure. 2021;29(4):385–92.

38. Bueno LL, Lobo FP, Morais CG, Mourão LC, de Ávila RAM, Soares IS, et al. Identification of a highly antigenic linear B cell epitope within Plasmodium vivax apical membrane antigen 1 (AMA-1). PloS one. 2011;6(6).

39. Temeeyasen G, Srijangwad A, Tripipat T, Tipsombatboon P, Piriyapongsa J, Phoolcharoen W, et al. Genetic diversity of ORF3 and spike genes of porcine epidemic diarrhea virus in Thailand. Infection, Genetics and Evolution. 2014;21:205–13.

40. Li Y, Ma ML, Lei Q, Wang F, Hong W, Lai DY, et al. Linear epitope landscape of the SARS-CoV-2 Spike protein constructed from 1,051 COVID-19 patients. Cell Rep. 2021;34(13):108915.

41. Li Y, Lai DY, Zhang HN, Jiang HW, Tian X, Ma ML, et al. Linear epitopes of SARS-CoV-2 spike protein elicit neutralizing antibodies in COVID-19 patients. Cell Mol Immunol. 2020;17(10):1095–7.

42. Shrock E, Fujimura E, Kula T, Timms RT, Lee IH, Leng Y, et al. Viral epitope profiling of COVID-19 patients reveals cross-reactivity and correlates of severity. Science. 2020;370(6520).

43. Shi J, Huang X, Liu Q, Huang Z. Identification of conserved neutralizing linear epitopes within the VP1 protein of coxsackievirus A16. Vaccine. 2013;31(17):2130–6.

44. Li C, Li W, de Esesarte EL, Guo H, van den Elzen P, Aarts E, et al. Cell attachment domains of the porcine epidemic diarrhea virus spike protein are key targets of neutralizing antibodies. Journal of virology. 2017;91(12):e00273–17.

45. Li W, van Kuppeveld FJ, He Q, Rottier PJ, Bosch B-J. Cellular entry of the porcine epidemic diarrhea virus. Virus research. 2016;226:117–27.

46. Deng F, Ye G, Liu Q, Navid M, Zhong X, Li Y, et al. Identification and comparison of receptor binding characteristics of the spike protein of two porcine epidemic diarrhea virus strains. Viruses. 2016;8(3):55.

47. Kirchdoerfer RN, Bhandari M, Martini O, Sewall LM, Bangaru S, Yoon KJ, et al. Structure and immune recognition of the porcine epidemic diarrhea virus spike protein. Structure. 2021;29(4):385–92 e5.

48. Chan W-E, Chuang C-K, Yeh S-H, Chang M-S, Chen SS-L. Functional characterization of heptad repeat 1 and 2 mutants of the spike protein of severe acute respiratory syndrome coronavirus. Journal of virology. 2006;80(7):3225–37.

49. Yi Z, Ling Y, Zhang X, Chen J, Hu K, Wang Y, et al. Functional mapping of B-cell linear epitopes of SARS-CoV-2 in COVID-19 convalescent population. Emerg Microbes Infect. 2020;9(1):1988–96.

50. Okonechnikov K, Golosova O, Fursov M. Unipro UGENE: a unified bioinformatics toolkit. Bioinformatics. 2012;28(8):1166–7.

51. Golosova O, Henderson R, Vaskin Y, Gabrielian A, Grekhov G, Nagarajan V, et al. Unipro UGENE NGS pipelines and components for variant calling, RNA-seq and ChIP-seq data analyses. PeerJ. 2014;2:e644.

52. Jespersen MC, Peters B, Nielsen M, Marcatili P. BepiPred-2.0: improving sequence-based B-cell epitope prediction using conformational epitopes. Nucleic Acids Research. 2017;45(W1):W24–W9.

53. Dosztanyi Z, Csizmok V, Tompa P, Simon I. The pairwise energy content estimated from amino acid composition discriminates between folded and intrinsically unstructured proteins. Journal of molecular biology. 2005;347(4):827–39.

54. Emini EA, Hughes JV, Perlow D, Boger J. Induction of hepatitis A virus-neutralizing antibody by a virus-specific synthetic peptide. Journal of virology. 1985;55(3):836–9.

55. Kolaskar A, Tongaonkar PC. A semi-empirical method for prediction of antigenic determinants on protein antigens. FEBS letters. 1990;276(1–2):172–4.

